# Improving detection capabilities of a critically endangered freshwater invertebrate with environmental DNA using digital droplet PCR

**DOI:** 10.1101/661447

**Authors:** Quentin Mauvisseau, John Davy-Bowker, Mark Bulling, Rein Brys, Sabrina Neyrinck, Christopher Troth, Michael Sweet

## Abstract

*Isogenus nubecula* is a critically endangered Plecoptera species. Considered extinct in the UK, the species was recently rediscovered in one location of the river Dee in Wales after 22 years of absence. As many species belonging to the Perlodidae, this species can be a bio-indicator, utilised for assessing water quality and health status of a given freshwater system. However, conventional monitoring of invertebrates via kick-sampling for example, is an invasive and expensive (time consuming). Further, such methods require a high level of taxonomic expertise. Here, we compared the traditional kick-sampling method with the use of eDNA detection using qPCR and ddPCR-analyses. In spring 2018, we sampled eDNA from twelve locations on the river Dee. *I. nubecula* was detected using kick-sampling in five of these locations, three locations using both eDNA detection and kick-sampling and one location using eDNA detection alone – resulting in a total of six known and distinct populations of this critically endangered species. Interestingly, despite the eDNA assay being validated in vitro and in silico, and results indicating high sensitivity, qPCR analysis of the eDNA samples proved to be ineffective. In contrast, ddPCR analyses resulted in a clear detection of *I. nubecula* at four locations suggesting that inhibition most likely explains the big discrepancy between the obtained qPCR and ddPCR results. It is therefore important to explore inhibition effects on any new eDNA assay. We also highlight that ddPCR may well be the best option for the detection of aquatic organisms which are either rare or likely to shed low levels of eDNA into their environment.

## Introduction

Monitoring biodiversity in freshwater systems is a cornerstone of the evaluation of the European Habitats Directive, the European Water Framework Directive and the general evaluation of ecosystem health and status ^1–3^. The assessment of freshwater biodiversity relies on biological monitoring methods, in which, the use of biodiversity indicators is an essential component of its evaluation. Various aquatic macroinvertebrates, such as mayflies, stoneflies and caddisflies (Ephemeroptera, Plecoptera and Trichoptera) are commonly used as bio-indicator organisms for water quality and ecosystem assessments ^4–6^. This is down to their rapid reaction to anthropogenic change such as levels of pollution, climate change, fracking, mining, and the construction of hydroelectric stations for example ^5,7–9^.

Traditional monitoring of macroinvertebrates via kick-sampling and/or capture-recapture methods, is, however, costly (i.e. time consuming), labour intensive and, above all, known to be limited in effective detection of populations below a certain threshold ^5,10^. Further, such methods are ecologically invasive i.e. they increase the risk of injury to the target (and non-target) organism. The morphological identification of these bio-indicators is also often challenging, especially at the immature life stages ^5,11–13^, so a high level of taxonomic expertise is therefore usually required to avoid any possible misidentification and therefore misrepresentation ^14,15^.

The use of molecular approaches for biomonitoring, for example the detection of environmental DNA (eDNA), may overcome a number of these issues, ^16^. Moreover, the use of eDNA increases efficiency, reliability and allows for a more rapid species identification and ultimately detection ^5^, whilst minimising any associated impacts on the species and the environment. All aquatic organisms shed DNA traces in their environment ^17^, and it is now possible to detect a specific species (barcoding) or assess an entire community (metabarcoding) by sampling an aquatic system and amplifying the existing DNA traces using PCR (Polymerase Chain Reaction) based techniques ^17^. Since the implementation of eDNA techniques in environmental studies last decade, it has been proven to be successful for the monitoring of invasive ^18–22^, endangered^23,24^ and/or economically important species from a wide range of taxa ^25–27^. However, few studies have used eDNA for monitoring rare or indicator macroinvertebrate species ^28–30^.

A typical example of such a bioindicator Plecoptera is the Scarce Yellow Sally stonefly, *Isogenus nubecula* (Perlodidae, Plecoptera) (Newman 1833). This critically endangered species has been reported as extinct or undetected in numerous countries from which it was originally present ^31,32^. Also in the UK, it was considered as extinct until recently, when *I. nubecula* specimens were rediscovered after a 22 year period of absence at a location in the river Dee, North Wales, UK ^32^. Moreover, a total of 14 individuals were recorded on that spot during two kick-sampling surveys. The aim of this study was to design a novel single species eDNA based primer/probe assay for the detection of *I. nubecula* and to compare the efficiency of qPCR (quantitative Polymerase Chain Reaction) and ddPCR (droplet digital Polymerase Chain Reaction) versus traditional kick-sampling

## Methods

### Primers and probe design

A species-specific set of primers and probe, targeting the COI gene (Cytochrome C Oxidase subunit 1 mitochondrial gene) of *I. nubecula* was designed using the Geneious Pro R10 Software (https://www.geneious.com; ^33^. This assay amplifies a 124 bp fragment using the forward primer (5’ – CCAGAAGCCTTGTAGAAAAC – 3’), the reverse primer (5’ – ACCCCGGCTAGATGAAGAGA – 3’) and a probe (6-FAM – CCCCACTCTCTGCTGGAATT – BHQ-1). Specificity of the assay was assessed *in-silico* by comparing against sequences from 21 genetically similar invertebrate species which had all been previously submitted to the NCBI (National Centre for Biotechnology Information; https://www.ncbi.nlm.nih.gov/) see Appendix 1 for full list. The specificity of the assay was tested *in-vitro* using PCR and qPCR, with DNA extracted from the nine invertebrate species (closely related or likely to be present in the same ecosystem). These included; *I. nubecula, Leuctra hippopus* (Kempny, 1899), *Perlodes mortoni* (Klapálek, 1906), *Nemoura lacustris* (Pictet, 1865), *Leuctra geniculata* (Stephens, 1836), *Nemoura erratica* (Claassen, 1936), *Taeniopteryx nebulosa* (Linnaeus, 1758), *Diura bicaudata* (Linnaeus, 1758) and *L. fusca* (Linnaeus, 1758).

### eDNA samples

12 locations from the River Dee, were sampled for eDNA between 9^th^ March 2018 and 1^st^ of April 2018 (Fig. 1 and Table 1). These locations were chosen following previous knowledge of historical observations in 1981 and 1982 ^32^. At each location, three independent (i.e. A, B and C) 1L water samples (referred to here after as natural replicates) were collected using a 40mL sterile polypropylene ladle and placed into a sterile plastic bag (Whirl-Pak^®^ 1242 ml Stand-Up Bag Merck^®^, Darmstadt, Germany) ^34^. Sub-samples were regularly collected from surface water downstream to upstream (to avoid disturbing sediments), across the width or the bank of the river, depending on the access and weather conditions following the method outlined in ^35^. Each independent 1L water sample was then filtered with a sterile 50 mL syringe (sterile Luer Lock™BD Plastipak™, Ireland) through a sterile 0.45 μm Sterivex™HV filter (Sterivex™filter unit, HV with luer-lock outlet, Merck^®^, Millipore^®^, Germany). Sterivex filters were immediately placed in a freezer bag and stored at – 80°C until further analysis in the laboratory. At each location, new sterile equipment and disposable nitrile gloves were used during the sampling process to avoid contamination. A ‘positive’ eDNA sample was collected by creating an isolated mesocosm onsite, which consisted of river water from site W4 and 11 specimens of *I. nubecula* stored for 1 hour. Two negative control samples were additionally filtered in the field with sterile ddH_2_O in parallel with the natural samples, to control for potential cross-contamination during the workflow.

**Table 1.** Table depicting the kick-sampling results for *I. nubecula* (i.e. how many specimens found at each site), the eDNA results using ddPCR analysis (i.e. if one natural replicate was positive to *I. nubecula* DNA), the amount of time spent performing kick-sampling and eDNA sampling, the amount of water filtrated for all natural replicate at each site, the sampling date, pH, dissolved oxygen and GPS coordinate. The site inaccessible for conducting a kick-sampling were marked “ns”.

| Sample ID | *I. nubecula* | eDNA (ddPCR) | Time (s) | Volume (ml) | Date | pH | O<sub>2</sub> | Latitude | Longitude |
| --- | --- | --- | --- | --- | --- | --- | --- | --- | --- |
| W1 | 0 | No | 60 | 350 | 31/03/2018 | 7.48 | 12.5 | 52.952759 | -3.0232733 |
| W2 | 3 | Yes | 60 | 200 | 01/04/2018 | 7.53 | 11.9 | 53.024980 | -2.8760059 |
| W3 | 30 | No | 60 | 700 | 09/03/2018 | 6.69 | 11.9 | 53.010679 | -2.8998019 |
| W4 | 16 | Yes | 120 | 1000 | 09/03/2018 | 6.52 | 11.8 | 53.003120 | -2.9138314 |
| W5 | ns | Yes | 45 | 750 | 15/03/2018 | 7.83 | 11.4 | 53.095257 | -2.8967275 |
| W6 | ns | No | 60 | 300 | 01/04/2018 | 7.82 | 12.5 | 53.011702 | -2.8686273 |
| W7 | 1 | Yes | 90 | 750 | 14/03/2018 | 7.67 | 11.6 | 52.978139 | -2.9627502 |
| W8 | 1 | No | 90 | 750 | 14/03/2018 | 7.8 | 10.7 | 52.964635 | -2.9628967 |
| W9 | 0 | No | 90 | 750 | 11/03/2018 | 6.75 | 11.8 | 52.945402 | -3.0194684 |
| W10 | 0 | No | 60 | 300 | 31/03/2018 | 7.74 | 13 | 52.970460 | -3.0879607 |
| W11 | 0 | No | 45 | 500 | 11/03/2018 | 6.63 | 11.6 | 53.100487 | -2.9239146 |
| W12 | 0 | No | 90 | 750 | 15/03/2018 | 7.69 | 10.9 | 52.967603 | -3.0619060 |

**Figure 1.**
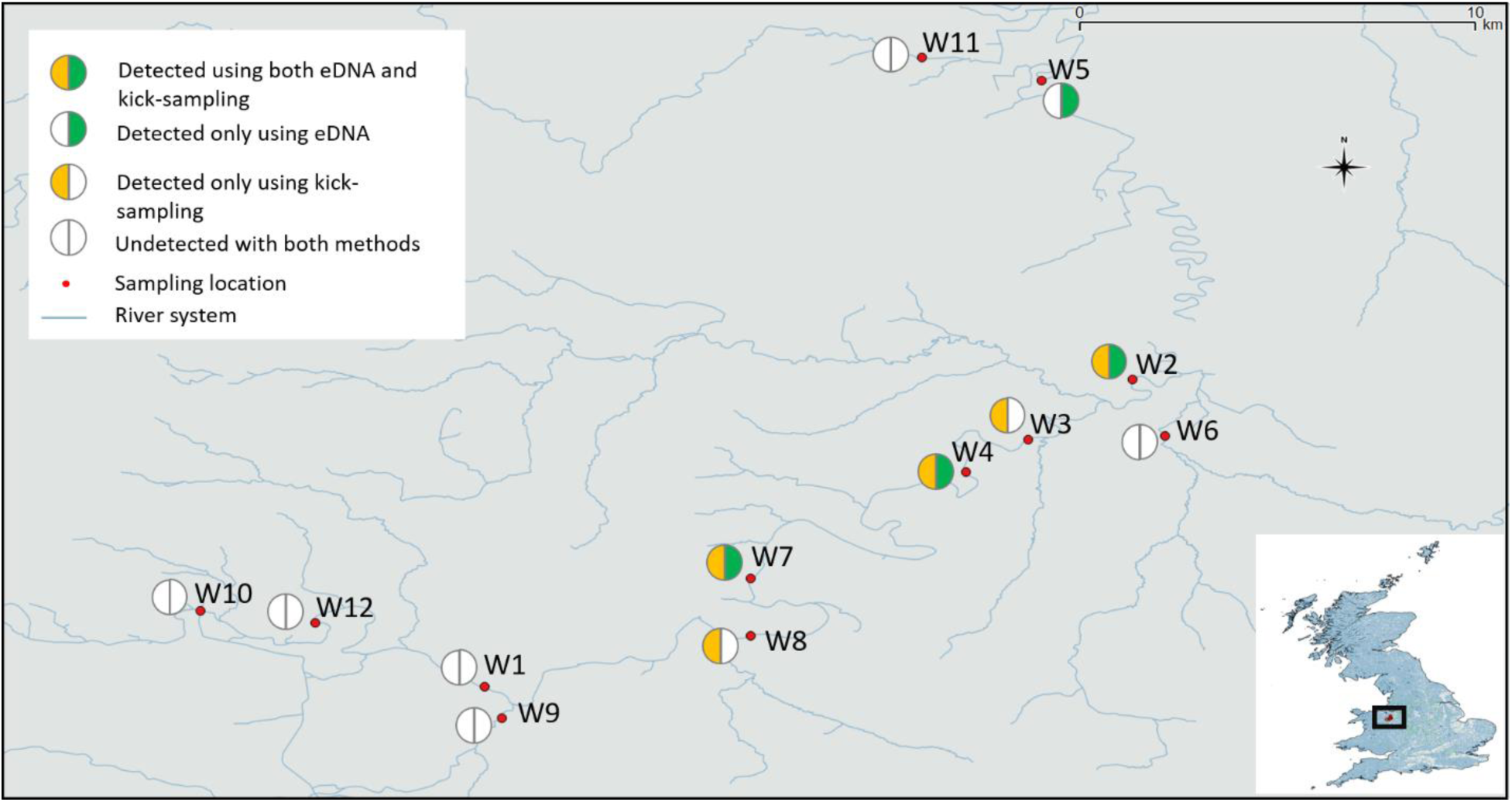
Map showing the 12 locations of the river Dee sampled with both kick-sampling and eDNA survey for monitoring I. nubecula in Wales, United Kingdom. Red dots are showing the sampled locations, half green circle the locations positive with eDNA detection using ddPCR, the half orange circle the locations were *I. nubecula* was found using kick-sampling. Locations W5 and W6 were not surveyed using kick-sampling method.

### DNA extraction

DNA extraction from both the eDNA samples and the tissue samples (utilised for validating the assay) was done using the Qiagen DNeasy^®^ Blood and Tissue Kit. We followed the manufacturer’s instructions for performing DNA extraction from tissue samples. Sterivex filters were extracted following the methods outlined in ^36^). All laboratory equipment was disinfected and decontaminated using UV-treatment prior to conducting any laboratory work. Laboratory equipment and surfaces were regularly disinfected using 10% bleach and absolute ethanol before conducting analyses.

### PCR

PCR amplifications were performed on a Gen Amp PCR System 9700 (Applied Biosystem) using the primers described above. PCR reactions were performed in a 25 µL reaction, with 12.5 µL of PCRBIO Ultra Mix Red (PCRBIOSYSTEMS), 1 µL of each primer (10 µM), 9.5 µL of ddH_2_O and 1 µL of template DNA. Optimal PCR conditions were performed under thermal cycling 50 °C for 2 min and 95 °C for 10 min, followed by 35 cycles of 95 °C for 15 s and 60°C for 1 min. For each PCR (with DNA from tissue samples), at least one positive and one negative control were included.

### qPCR

qPCR amplifications were performed on an ABI StepOnePlus™ Real-Time PCR (Applied Biosystems) in final volumes of 25 µL, using 12.5 µL of PrecisionPlus qPCR Master Mix with ROX (Primer Design, UK), 1 µL of each primer (10 µM), 1 µL of probe (2.5 µM), 6.5 µL of ddH_2_O and 3 µL of extracted DNA. qPCR conditions were as follow: 50 °C for 2 min and 95 °C for 10 min, followed by 45 cycles of 95 °C for 15 s and 60 °C for 1 min. For each qPCR with DNA from tissue samples, at least two positive and two negative controls were included. A standard curve was established by analysing a 1:10 dilution series of DNA extracted from *I. nubecula* (68.2 ng/ µL, Nanodrop 2000 Spectrophotometer, ThermoFisher Scientific) following the MIQE Guidelines ^37^ (Appendix 2).

### ddPCR

Digital droplet PCR was conducted using the Bio-Rad QX200 ddPCR system in a 20-μl total volume. Each reaction contained 10 μL Bio-Rad ddPCR supermix for probes (no dUTP), 750 nM of each primer, 375 nM probe, 3 µL DEPC water, and 4 µl template DNA. Twenty microlitres of the PCR mix was pipetted into the sample chambers of a Droplet Generator DG8 Cartridge (Bio-Rad, cat no. 1864008), and 70 μL of the Droplet Generation Oil for Probes (Bio-Rad, cat no. 186-4005) was added to the appropriate wells. The cartridges were covered with DG8 Gaskets (Bio-Rad, cat no. 1863009) and placed in a QX200 Droplet Generator (Bio-Rad) to generate the droplets. After droplet generation, the droplets (40 μL) were carefully transferred to a ddPCR 96-well plate (Bio-Rad, cat no. 12001925). The PCR plate was sealed with pierceable foil (Bio-Rad, cat no. 181-4040). PCRs were performed using a C1000 Touch™ Thermal Cycler with a 96-well Deep Reaction Module (Bio-Rad). PCR conditions were 10 min at 95°C, followed by 40 cycles of denaturation for 30 s at 94°C and extension at 60°C for 1 min, with ramp rate of 2°C s-1, followed by 10 min at 98°C and a hold at 12°C. Droplets were then read on a QX200 droplet reader (Bio-Rad). All droplets were checked for fluorescence and the Bio-Rad’s QuantaSoft software version 1.7.4.0917 was used to quantify the number of *I. nubecula* copies per µL. Thresholds for positive signals were determined according to QuantaSoft software instructions. All droplets beyond the fluorescence threshold (3500) were counted as positive events, and those below it as negative events. All eDNA samples were analysed in duplicate (one replicate undiluted and one replicate diluted 1:2). One positive control (i.e. DNA extracted from *I. nubecula* at a concentration of 1 ng/ µL diluted 1:100 (Nanodrop 2000 Spectrophotometer, ThermoFisher Scientific)), one No Template Control (i.e., IDTE pH 5.0) and the two negative field controls were additionally included. The LOD using the ddPCR was assessed following the method outlined in ^38^. We conducted a serial dilution of a DNA extracted from *I. nubecula*. The starting point was an initial 1: 100 dilution of extracted genomic DNA from *I. nubecula* at 1 ng/ µL, followed by a serial 1:5 dilution. The serial dilution included ten replicate of each dilution.

### Estimation of the LOD and LOQ

To become estimates of the LOD and LOQ for the primer/probe assay used on both the qPCR and ddPCR machines, we set-up a dilutions range from 10^-1^ to 10^-9^ with 10 technical replicates used for each of the dilution steps. Following ^37^, the LOD was defined as the lowest concentration in which 95% of positive samples were detected. The LOQ was defined as the last standard dilution in which the targeted DNA was detected and quantified in at least 90% of positive samples ^34,35^. All eDNA samples were then analysed with six technical replicates ^34,35^ on a qPCR plate, with six negative controls and a positive control dilution series from 10^-1^ to 10^-6^ in duplicate.

### Kick-sampling

Kick-sampling was performed using the standard of the Freshwater Biological Association (UK), i.e. using a kick-sampling net with a 1 mm mesh (see detailed protocol: https://www.fba.org.uk/practical-guidance-sampling-and-collecting). Sampling duration was recorded at each site and varied depending on access, depth, river flow, or weather conditions (Table 1). Perlodidae specimens found during kick-sampling were either preserved in 99% ethanol or kept alive as a part or a separate rearing experiment. Specimens were identified in the laboratory by two independent taxonomy experts (John Davy-Bowker & Michael Hammett) using a low-power binocular microscope with cold light source and using an identification key ^39,40^.

### Statistical analysis

A site occupancy modelling approach ^41–43^ was utilised to assess the effect of environmental covariates on the presence of eDNA of *I. nubecula* and to estimate the detection probability. This hierarchical modelling framework has the advantage of accounting for the risk of false negative results when estimating the probability of detection. This analysis was run with the ddPCR data (Appendix 3). Covariates tested included: (i) turbidity (likely to inhibit the PCR reaction, with the volume of filtered water being used as a proxy), (ii) pH, (iii) dissolved oxygen concentration, (iv) amount of time including eDNA sampling and kick-sampling spent at each location as indicator of the field conditions and (v) human accessibility as a binary indicator (possible to perform kick-sampling/absence of kick-sampling survey) (Appendix 4). Analyses were performed using the ‘eDNAoccupancy’ package ^44,45^ in the R statistical programming environment (R Core Team, 2018). Model selection and interpretation followed procedures given in ^44,45^. We fitted our model using the ‘occModel’ function from the described package. MCMC chains ran for 11,000 iterations, with 10,000 retained for obtaining parameter estimates and credible intervals.

## Results

### Specificity and validation of eDNA assay using PCR, qPCR and ddPCR

The primers and probe designed in this study were species-specific *in-silico* and *in-vitro* with both conventional PCR and qPCR. The negative controls or samples with DNA from non-target species did not amplify with either method. For qPCR, we analysed the standard curve and compiled the limit of detection (LOD) and limit of quantification (LOQ) as per the MIQE guidelines ^34,37^. The LOD was 6.82 × 10^6^ng DNA µL^-1^ at 39.29 ± 2.00 Ct (i.e. Cycle threshold) and the LOQ was 6.82 × 10^-4^ ng DNA µL^-1^ at 34.48 ±0.95 Ct (Slope= −3.86, Y inter= 19.52, R^2^= 0.97, Eff%= 81.63) (Figure 2). Using ddPCR, five replicates from the dilution which equated to 0.08 pg of DNA yielded a positive detection (mean 0.05 copy per µL^-1^) and only one replicate of the next dilution (i.e. 0.016 pg) yielded to a positive detection of *I. nubecula* (0.08 copy per µL^-1^). All replicates from further dilution and negative controls were negative. However, as specified in ^38^ (at the lower end of detection), the lower 95% confidence can limit overlap with potential artefact in the negative control. For this reason, we considered 0.08 pg of DNA to be the lowest amount able to be detected using ddPCR and only considered samples > 0.5 copy per µL^-1^ (as in ^38^) to meet the threshold for a positive detection.

**Figure 2.**
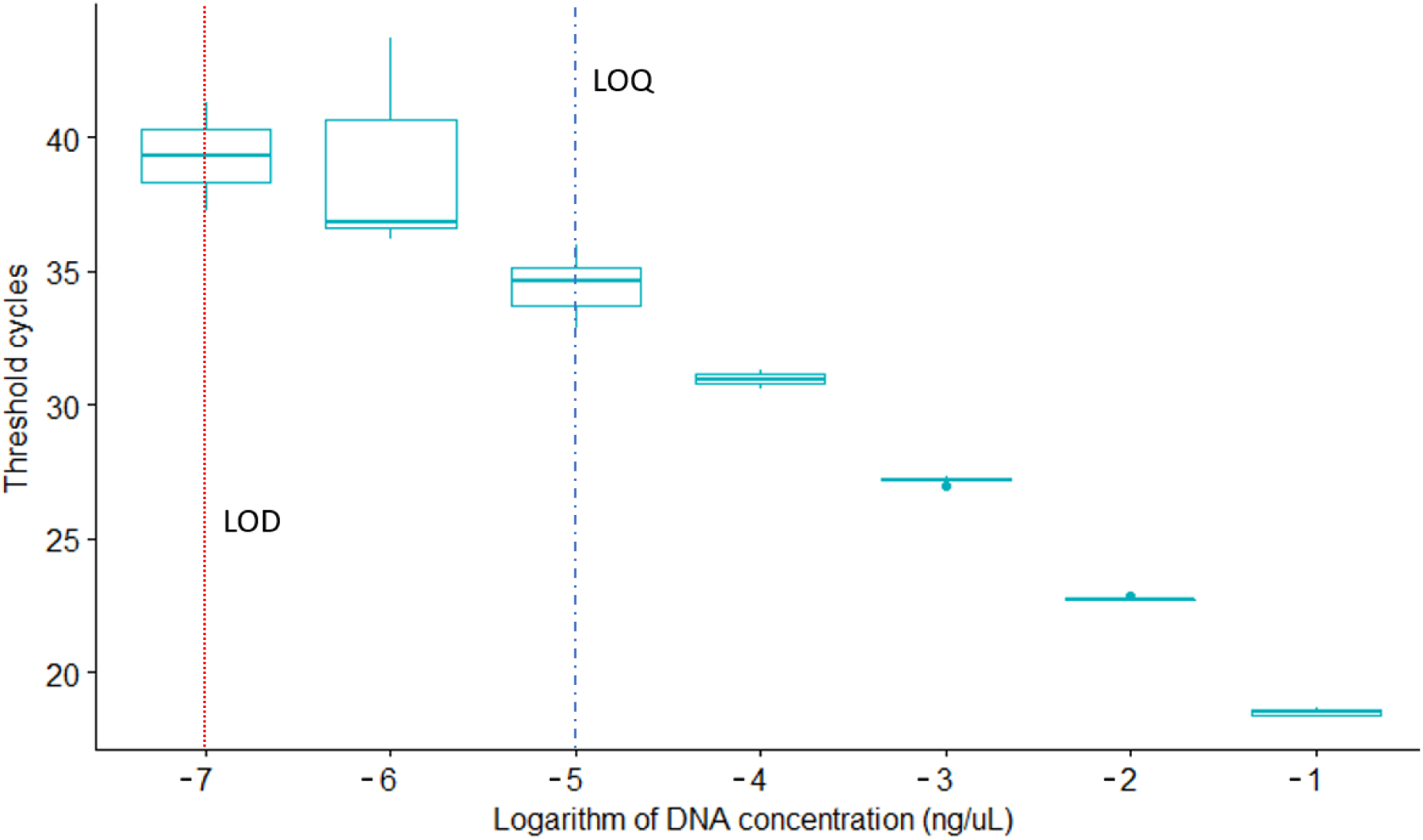
Standard curve assessing the Limit of Detection (LOD) and Limit of Quantification (LOQ) for the qPCR assays detecting the DNA traces of *I. nubecula*. Both limits were calculated from a 1:10 serial dilution with 10 replicates per concentration. The LOD was 6.82 10-6 ng DNA µL-1 at 39.29 ± 2.00 Ct (i.e. Cycle threshold) and the LOQ was 6.82 10-4 ng DNA µL-1 at 34,48 ± 0,95 Ct (Slope= −3.86, Y inter= 19.52, R^2^= 0.97, Eff%= 81.63).

### Kick sampling assessment

*I. nubecula* was found at 5 sites along the River Dee, whereas the species could not been found at five other sites (Fig. 1, Table 1). Abundance ranged from just 1 individual at two sites, at W7 and W8, up to a highest density of 30 individuals at W3. Two of the sites surveyed for eDNA were not assessed via kick sampling due to dangerous access and weather conditions (Table 1).

### Comparison of qPCR versus ddPCR analyses

Despite the success of the assay *in-silico* and *in-vitro*, no amplification could be obtained via qPCR on each of the eDNA samples (Table 2). Even the ‘positive control eDNA sample’ which consisted of 11 *I. nubecula* individuals housed in a 1 litre mesocosm for a period of one hour before filtering (see methods). During each run, the positive dilution range indicated the assay ran without any issue (Slope= −3.65 / −4.05, Y inter= 19.22 / 26.46, R^2^= 0.98 / 0.99, Eff%=76.46 / 88.03). In contrast, the ddPCR analysis revealed a positive detection of *I. nubecula* at four sampling locations (Figure 1, Table 2). Concentration ranged from 0.6 to 0.14 copy per µL^-1^ in the eDNA samples. The ‘positive eDNA’ sample generated a DNA concentration of 5.4 copies per µL^-1^ (undiluted) and 8.2 copies per µL^-1^ (diluted).

**Table 2.**
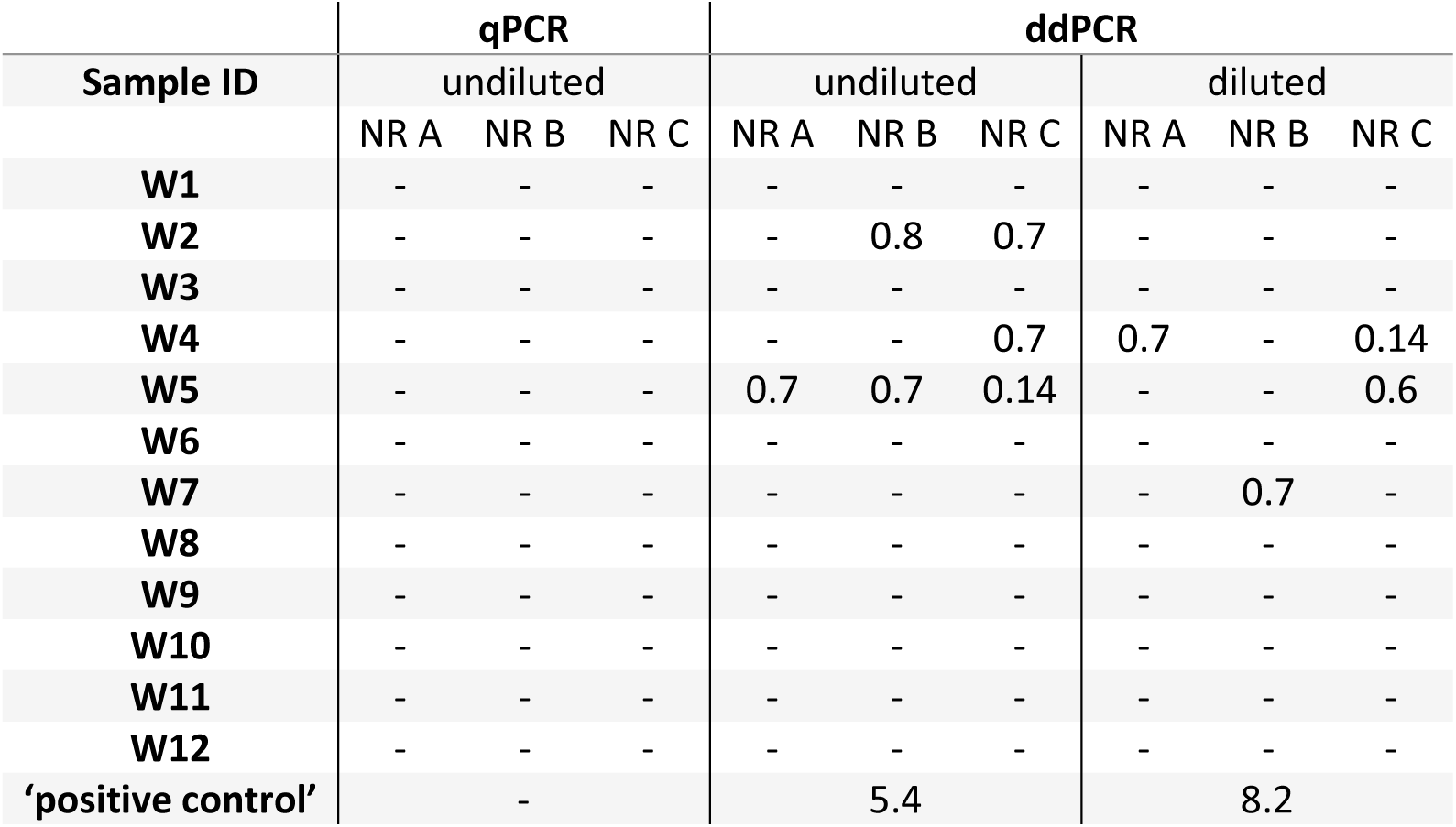
Table depicting the eDNA detection results using qPCR and ddPCR techniques on diluted and undiluted (1:2) natural replicates (NR) sampled at each field location. ‘-’ depict the absence of eDNA detection using qPCR and/or ddPCR. Quantification values of ddPCR results are displayed in copy per µL^-1^. Natural replicates were analysed using six technical replicates with qPCR and without replicates using ddPCR. All samples revealed a negative result for *I. nubecula* eDNA using qPCR. DNA from the targeted specie was amplified in samples from four field locations and in the ‘positive control’.

The site occupancy modelling approach did not reveal any significant effect of the environmental variables on the presence of eDNA or on the probability of detection (Tables 3, 4). Probabilities of *I. nubecula* occurrence were relatively low and ranging from 0.45 to 0.53 (Table 4). Probabilities of eDNA detection at each sampling sites ranged from 0.59 at site W5 where all ‘natural replicates’ where found to be positive using ddPCR to 0.27 at site W10, a site with high turbidity where no stonefly were found.

**Table 3.**
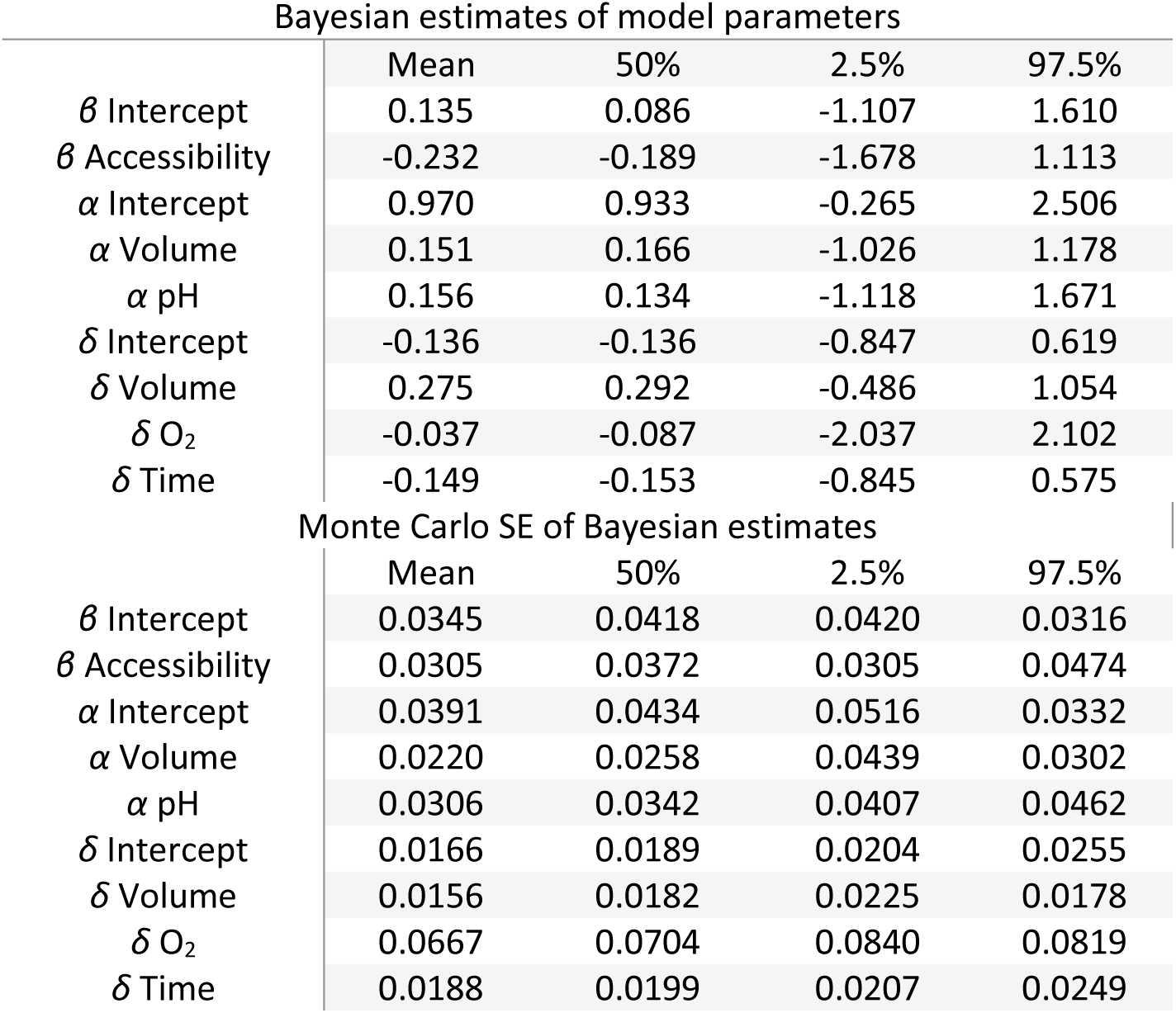
Table depicting the of the Bayesian estimates for effects of covariates on the probability of occurrence at a site (*ψ*). (*α*) and (*δ*) parameters are covariates for the conditional probability of eDNA presence in a sample (θ) and for its detection (*p*). (*ß*) parameters are covariates of the estimated occupancy (*ψ*). Means represent estimated parameter values and last two columns represent the boundaries of the 95% credible intervals.

**Table 4.** Table depicting the Bayesian estimates for the probabilities of occurrence (*ψ*), the conditional probabilities of eDNA presence in a sample (ϑ) and eDNA detection (*p*) of *I. nubecula* at each sampling site of the river Dee and its tributaries.

| Site | $\psi$ | $\vartheta$ | $p$ |
| --- | --- | --- | --- |
| W1 | 0.45 | 0.79 | 0.33 |
| W2 | 0.45 | 0.76 | 0.31 |
| W3 | 0.45 | 0.77 | 0.52 |
| W4 | 0.45 | 0.75 | 0.48 |
| W5 | 0.53 | 0.87 | 0.59 |
| W6 | 0.53 | 0.81 | 0.32 |
| W7 | 0.45 | 0.86 | 0.46 |
| W8 | 0.45 | 0.87 | 0.52 |
| W9 | 0.45 | 0.77 | 0.47 |
| W10 | 0.45 | 0.80 | 0.27 |
| W11 | 0.45 | 0.75 | 0.49 |
| W12 | 0.45 | 0.86 | 0.50 |

## Discussion

In our study, we compared the use of kick-sampling and eDNA detection for monitoring a critically endangered bioindicator macroinvertebrate. While our eDNA detection approach using qPCR showed high sensitivity (Figure. 2), with no false positive results during the validation process and assessment of the MIQE guidelines (Appendix 2), we were, however, not able to amplify DNA traces of *I. nubecula* in any of the eDNA samples. This is surprising as one should expect positive detection at least in the five locations where we found the species via kick-sampling, and especially in the ‘positive eDNA’ sample. These observations thus clearly pose doubts on the concept of eDNA using the qPCR methodology.

Potential explanations for these false negative observations might be (i) an incorrect sampling protocol, (ii) the presence of PCR inhibitors in the DNA extracts, or (iii) a very limited shedding rate of the targeted species ^46^. As previously shown, the sampling design of any eDNA based study can affect the reliability of detection ^34^. In this case, however, we accounted for this by taking, for example, three natural replicates at each site and incorporating six technical PCR replicates per sample.

Most likely, inhibition of the qPCR assay is the most responsible aspect for the false negative detections, is it has also been found to be the case in other studies ^47,48^. One can assess for inhibition via the use of internal positive controls, such as spiked synthetic DNA or different from the targeted species ^46^. Limited detection or complete failure of such internal controls may then clearly show the occurrence of inhibition factors. If there is inhibition, two methods can be used to overcome this issue. The first method is to dilute the DNA extracted from the field sample ^46^, whilst the second is the use of an inhibitor removal kit ^46,49^. However, both methods have been shown to reduce the yield of target DNA in the extracted sample ^46^. In our study, qPCR showed no results from the eDNA samples and so we hypothesised that inhibition may be an important driver for the false negative observations in this assay. We did not use an inhibitor removal kit in order to avoid reducing the amount of DNA extracted from the field samples. Instead, we ran the samples on a ddPCR with two different dilutions. ddPCR has been shown to outperform qPCR in some other studies ^45,50–52^ by simply detecting and quantifying lower amounts of DNA, and being less sensitive to inhibition. Our findings also support these observations as we were able to detect the presence of *I. nubecula* eDNA at four distinct locations. Three of them matched with the positive results from the kick-sampling survey. Interestingly, the analysis of the ‘positive eDNA sample’ showed an increase from 5.4 copies per µL (undiluted) to 8.2 copies per µL (diluted), indicating that inhibition was still affecting the ddPCR (although not strong enough to block amplification in this instance). This finding indicates that there are substantial inhibiting factors affecting the primer/probe assay used and may explain the false negative results following qPCR.

The very low *I. nubecula* eDNA concentrations in the samples also indicate that this species is characterized by very low shedding rates. Moreover, in all locations, the eDNA concentration ranged from only 0.6 to 0.14 copies per µL and up to 8.2 copies per µL in the ‘positive eDNA’ sample. As this study is the first to use ddPCR for detecting low populations of endangered invertebrates in fast flowing rivers, we cannot compare our results with previously published studies. Besides the fact that invertebrates are generally found to shed only limited amounts of eDNA in the water, potential other explanatory variables could be the high flow rate of the river and low temperature during sampling. Sampling was undertaken at the end of winter/beginning of spring, when environmental conditions such as high flow rates or flood events could have decreased and diluted the quantity of DNA traces. However, this was unavoidable for this species as *I. nubecula* emerges from March onwards ^31,39^ and so for this species sampling time could not be altered.

Finally, when sampling for any eDNA study, it is useful to have an understanding the ecology of the species under study, such as the species habits and preferred habitat in which it occurs. However, again, as *I. nubecula* was recently rediscovered in Wales, there is very little information on this species ^32^. Our site occupancy modelling approach was also not able to identify any specific variable which would have a significant effect on the probability of detection of this species (Figure 3, Tables 3, 4), which is quite logic as all the sites were located in the same study system. A recent study by ^45^ on Burmese pythons similarly acknowledge that occupancy modelling approach analyses have certain limitations, mainly driven by the number of locations sampled and restricted range of environmental values collected. In addition, the species in question appears to be rare, and its distribution may be subject to high degrees of stochasticity with regard to population dynamics. Thereby resulting in the effects of the underlying environmental drivers of its distribution being harder to detect. Further work will therefore be necessary in order to increase our understanding of the ecology of *I. nubecula* if we want to optimize the sampling protocol and conservation plans for this species. Notably, site occupancy modelling is most flexible using the Bayesian statistical framework, and this allows the combination of prior information along with information gained from new sampling data to produce a more informed post experimental understanding, allowing the combination of previous data with current data to produce more robust results ^43,53^.

**Figure 3.**
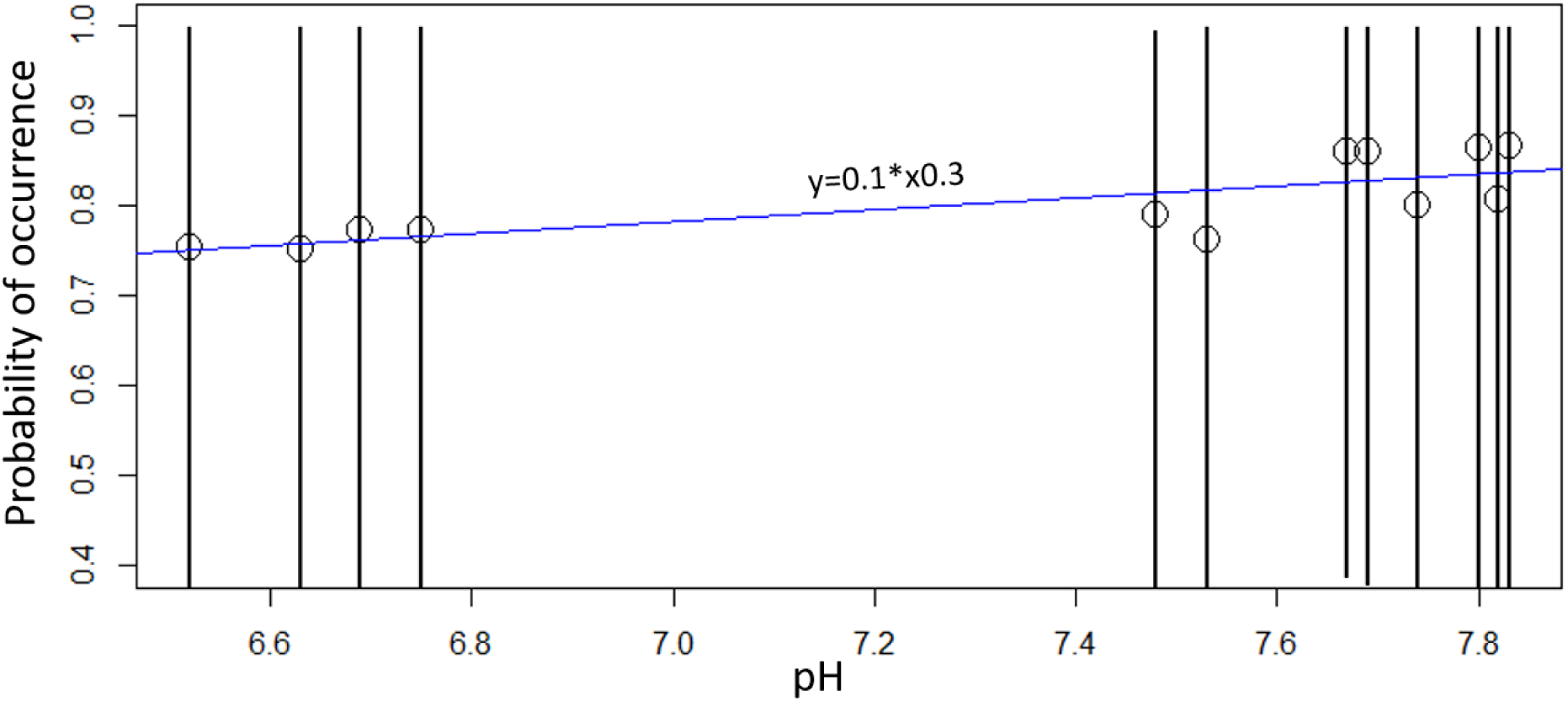
Estimated probability of occurrence of *I. nubecula* eDNA with the pH of each sampling sites. Dots are representing each sampling locations, the black lines are representing the estimates of posterior medians with 95% credible intervals and the blue line the regression analysis.

In conclusion, even if the highest standards of validation are undertaken in the design and implementation of an eDNA based PCR or qPCR assay ^28–30^, false negative results can appear by inhibition factors ^46^, low shedding rates from the target species ^18,54^ or low population sizes ^20^. In this case we are dealing with extreme scenario, in which none of our eDNA samples showed any amplification via qPCR despite the fact that populations of *I. nubecula* were present. However, we got positive detection (using ddPCR) at most of the locations where the species was found via the physical survey effort. Less than ten studies have (at the time of writing) utilised this technology for eDNA assays ^38,45,50–52,55–57^, but this is likely to increase significantly due to the apparent benefits observed in this study for example. Caution should therefore be taken with any negative results derived from assays reliant solely on qPCR for the reasons given above.

## Acknowledgement

We would like to thank Mike Howe from Natural Resources wales and Michael Hammett for helping during the stonefly identification process. The study was funded by Surescreen Scientifics, UK and Natural Resources Wales.

## Author contributions

Q.M. and M.S. designed the experiment and methodology; Q.M., M.S., C.T and J.D.B collected field samples; Q.M. performed extraction and qPCR; S.N performed ddPCR; Q.M. analysed the data. The manuscript was written by Q.M, M.B and M.S and reviewed by all authors.

## Supplementary information

Appendix 1. List of invertebrate species and the related GenBank accession number

Appendix 2. MIQE Guidelines *Isogenus nubecula*

Appendix 3. Detection Data *Isogenus nubecula*

Appendix 4. Survey Data *Isogenus nubecula*

