## Appendices 1-4 for "Improving detection capabilities of a critically endangered freshwater invertebrate with environmental DNA using digital droplet PCR"

Appendix 1: List of invertebrate species and the related GenBank accession number utilized when developing and validating the species-specific primers and probe used in this study.

| **Species** | **Accession number** |
|---|---|
| *Isogenus nubecula* (Newman, 1833) | MF801622.1 |
| *Amphinemura standfussi* (Ris, 1902) | JX460920 |
| *Amphinemura sulcicollis* (Stephens, 1836) | JX495637 |
| *Brachyptera risi* (Morton, 1896) | KF492801 |
| *Capnia atra* (Morton, 1896) | KF809153 |
| *Zwicknia bifrons* (Newman, 1838) | KF144842 |
| *Capnia vidua* (Klapálek, 1904) | JQ736348 |
| *Chloroperla tripunctata* (Scopoli, 1763) | HQ705654 |
| *Cotesia acuminate* (Reinhard, 1880) | AY333870 |
| *Dinocras cephalotes* (Curtis, 1827) | KF492802 |
| *Diura bicaudata* (Linnaeus, 1758) | KJ675053.1 |
| *Heptagenia longicauda* (Stephens, 1836) | LN734744 |
| *Isoperla grammatica* (Poda, 1761) | KU955895 |
| *Isoperla obscura* (Zetterstedt, 1840) | KJ675043 |
| *Kageronia fuscogrisea* (Retzius, 1783) | JN299122 |
| *Leuctra fusca* (Linnaeus, 1758) | KT807840.1 |
| *Leuctra hippopus* (Kempny, 1899) | KF809176.1 |
| *Nemoura avicularis* (Morton, 1894) | JX905857 |
| *Nemoura cinerea* (Retzius, 1783) | JX495661 |
| *Nemurella pictetii* (Klapálek, 1900) | KF492804 |
| *Protonemura meyeri* (Pictet, 1841) | KF492803 |
| *Sterrhopterix standfussi* (Wocke, 1851) | HM873931 |
| *Nemoura lacustris* (Pictet, 1865) | MF801623.1 |

### Appendix 2. MIQE Guidelines (*Isogenus nubecula* )

| ITEM TO CHECK | IMPORTANCE | CHECKLIST |
|---|---|---|
| **EXPERIMENTAL DESIGN** | | COI |
| Definition of experimental and control groups | E | 12 locations from the River Dee in Wales, UK, were sampled between |
| Number within each group | E | 12 locations from the River Dee in Wales, UK, were sampled between |
| Assay carried out by core lab or investigator's lab? | D | Investigator's Lab |
| Acknowledgement of authors' | D | Yes |
| SAMPLE | | |
| Description | E | Field samples |
| Volume/mass of sample processed | D | Three independant water samples ranging from 200 ml to 1000 ml were |
| Microdissection or | E | N/A |
| Processing procedure | E | Each water sample (i.e. natural replicates) were filtered using a sterile 0.45 µm Sterivex™ HV filter (Sterivex™ filter unit, HV with luer-lock outlet, Merck®, Millipore®, Germany). The filters were stored in a freezing bag with several pack of ice on site and then stored at –80 °C until DNA extraction. DNA was extracted using Qiagen® DNA extraction Kit ( DNeasy Blood & Tissue Kits) as per Spens et al. 2017.The final volume of an extracted sample was 100 µL with Buffer AE and stored at –20 °C until qPCR. |
| If frozen - how and how quickly? | E | The filters were stored in a freezing bag with several pack of ice on site and then stored at –80 °C until DNA extraction. |
| If fixed - with what, how quickly? | E | |
| Sample storage conditions and duration (especially for FFPE samples) | E | All concentrated samples were stored in 1.5 mL eppendorf tubes at –20°C. |
| NUCLEIC ACID EXTRACTION | | |
| Procedure and/or instrumentation | E | We used DNA extraction kit from Qiagen DNeasy blood and tissue kit |

| | | |
|---|---|---|
| Name of kit and details of any modifications | E | We used DNA extraction kit from Qiagen DNeasy blood and tissue kit as per Spens et al. 2017 |
| Source of additional reagents used | D | N/A |
| Details of DNase or RNAse treatment | E | N/A |
| Contamination assessment (DNA or RNA) | E | Two "Blank" filters with filtrated ddH2O were extracted with the eDNA samples from the mesocosm experiment. |
| Nucleic acid quantification | E | Quantification for the standard curve was performed using a Nanodrop 2000 Spectrophotometer, (Thermofisher Scientific) following the manufacturer's instructions. |
| Instrument and method | E | |
| Purity (A260/A280) | D | |
| Yield | D | |
| RNA integrity method/instrument | E | N/A |
| RIN/RQI or Cq of 3' and 5' transcripts | E | N/A |
| Electrophoresis traces | D | N/A |
| Inhibition testing (Cq dilutions, spike or other) | E | Not Checked |
| REVERSE TRANSCRIPTION | | |
| Complete reaction conditions | E | N/A |
| Amount of RNA and reaction volume | E | N/A |
| Priming oligonucleotide (if using GSP) and concentration | E | N/A |
| Reverse transcriptase and concentration | E | N/A |
| Temperature and time | E | N/A |
| Manufacturer of reagents and catalogue numbers | D | N/A |
| Cqs with and without RT | D | N/A |
| Storage conditions of cDNA | D | N/A |
| qPCR TARGET INFORMATION | | |
| If multiplex, efficiency and LOD of each assay. | E | N/A |
| Sequence accession number | E | N/A |
| Location of amplicon | D | N/A |
| Amplicon length | E | 124 bp fragment (both primers included) |

| | | |
|---|---|---|
| In silico specificity screen (BLAST, etc) | E | Primers and probes were found to be specific in silico using NCBI website (https://www.ncbi.nlm.nih.gov/) and Geneious Pro R10 software. |
| Pseudogenes, retropseudogenes or other homologs? | D | Not Found |
| Sequence alignment | D | N/A |
| Secondary structure analysis of amplicon | D | Not Checked |
| Location of each primer by exon or intron (if applicable) | E | N/A |
| What splice variants are targeted? | E | N/A |
| qPCR OLIGONUCLEOTIDES | | |
| Primer sequences | E | Forward primer (5' – CCAGAAGCCTTGTAGAAAAC – 3') Reverse primer (5' – ACCCCGGCTAGATGAAGAGA – 3') |
| RTPrimerDB Identification Number | D | Not Submitted |
| Probe sequences | D | Probe (6-FAM – CCCCACTCTCTGCTGGAATT – BHQ-1) |
| Location and identity of any modifications | E | |
| Manufacturer of oligonucleotides | D | (Sigma-Aldricht) Merck KGaA, Darmstadt, Germany |
| Purification method | D | HPLC |
| qPCR PROTOCOL | | |
| Complete reaction conditions | E | Reactions were set up manually in a specific cabinet using designated equipment. |
| Reaction volume and amount of cDNA/DNA | E | Reaction volume is 25 µL, and amount of DNA is 3 µL |
| Primer, (probe), Mg++ and dNTP concentrations | E | qPCR amplification was performed in a final volume of 25 µl using 12.5 µl of PrecisionPlus qPCR Master Mix with ROX (Primer Design, UK), 1 µl of each primer (10 µM), 1 µl of the corresponding probe (2.5 µM), 6.5 µl of ddH2O and 3µl of extracted DNA |
| Polymerase identity and concentration | E | We used PrecisionPlus qPCR Master Mix with ROX (Primer Design, UK) |
| Buffer/kit identity and manufacturer | E | We used PrecisionPlus qPCR Master Mix with ROX (Primer Design, UK) |
| Exact chemical constitution of the buffer | D | N/A |

| | | |
|---|---|---|
| Additives (SYBR Green I, DMSO, etc.) | E | N/A |
| Manufacturer of plates/tubes and catalog number | D | MicroAmp® Fast 96-Well Reaction plate (0,1ml) Applied Biosystems, Warrington, UK with optical adhesive covers Applied Biosystems, Warrington, UK |
| Complete thermocycling parameters | E | warm up at 50°C for 2 min and denaturation at 95°C for 10 min, followed by 45 cycles 95°C for 15s, 60°C for 1 min |
| Reaction setup (manual/robotic) | D | We performed following the manufacturer's instructions. |
| Manufacturer of qPCR instrument | E | ABI StepOnePlus™ Real-Time PCR (Applied Biosystems, Warrington, UK). |
| qPCR VALIDATION | | |
| Evidence of optimisation (from gradients) | D | qPCR conditions were found with an anealing temperature of 60°C |
| Specificity (gel, sequence, melt, or digest) | E | The specificity of the primers-probe targeting the COI were tested against DNA extracted from : I. nubecula, Leuctra hippopus (Kempny 1899), Perlodes mortoni (Klapálek 1906), Nemoura lacustris (Pictet 1865), Leuctra geniculata (Stephens 1836), Nemoura erratica (Claassen 1936), Taeniopteryx nebulosa (Linnaeus 1758), Diura bicaudata (Linnaeus 1758) and Leuctra fusca (Linnaeus 1758). Primers/probe designed in this study were found to be species-specific to Isogenus nubecula. |
| For SYBR Green I, Cq of the NTC | E | N/A |
| Standard curves with slope and y-intercept | E | Slope= -3.858, Y inter= 19.52 |
| PCR efficiency calculated from slope | E | Eff%= 81.625 |
| Confidence interval for PCR efficiency or standard error | D | Not Checked |
| r2 of standard curve | E | $R^2$= 0.966 |
| Linear dynamic range | E | Not Checked |
| Cq variation at lower limit | E | The positive signals were detected from at least one of the six qPCR replicates per eDNA sample |

| | | |
|---|---|---|
| Confidence intervals throughout range | D | Not Checked |
| Evidence for limit of detection | E | Based on the standard |
| If multiplex, efficiency and LOD of each assay. | E | N/A |
| DATA ANALYSIS | | |
| qPCR analysis program (source, version) | E | StepOnePlus™ Software v2.2.2 |
| Cq method determination | E | We performed according to default setting of Software above. |
| Outlier identification and disposition | E | N/A |
| Results of NTCs | E | At least six wells of no-template negative control were adopted in all qPCR plates |
| Justification of number and choice of reference genes | E | N/A |
| Description of normalisation method | E | We used standard curve methods. |
| Number and concordance of biological replicates | D | N/A |
| Number and stage (RT or qPCR) of technical replicates | E | We performed in 6 replicate for each eDNA samples. We performed the standard curve with 10 replicates for each dilution. |
| Repeatability (intra-assay variation) | E | Not Checked |
| Reproducibility (inter-assay variation, %CV) | D | Not Checked |
| Power analysis | D | N/A |
| Statistical methods for result significance | E | We performed according to default setting of Software above. |
| Software (source, version) | E | StepOnePlus™ Software v2.2.2 |
| Cq or raw data submission using RDML | D | Not Submitted |

Information (Bustin et al. 2009). (E) means Essential information and (D) means Desirable information.

Bustin, S. A., Benes, V., Garson, J. A., Hellemans, J., Huggett, J., Kubista, M., … Wittwer, C. T. (2009). The MIQE Guidelines: Minimum Information for Publication of Quantitative Real-Time PCR Experiments. Clinical Chemistry, 55(4), 611–622. doi:10.1373/clinchem.2008.112797

Appendix 3. Detection Data *Isogenus nubecula*

| site | sample | pcr1 | pcr2 |
|---|---|---|---|
| W1 | 1 | 0 | 0 |
| W2 | 1 | 0 | 0 |
| W3 | 1 | 0 | 0 |
| W4 | 1 | 0 | 1 |
| W5 | 1 | 1 | 0 |
| W6 | 1 | 0 | 0 |
| W7 | 1 | 0 | 0 |
| W8 | 1 | 0 | 0 |
| W9 | 1 | 0 | 0 |
| W10 | 1 | 0 | 0 |
| W11 | 1 | 0 | 0 |
| W12 | 1 | 0 | 0 |
| W1 | 2 | 0 | 0 |
| W2 | 2 | 1 | 0 |
| W3 | 2 | 0 | 0 |
| W4 | 2 | 0 | 0 |
| W5 | 2 | 1 | 0 |
| W6 | 2 | 0 | 0 |
| W7 | 2 | 0 | 1 |
| W8 | 2 | 0 | 0 |
| W9 | 2 | 0 | 0 |
| W10 | 2 | 0 | 0 |
| W11 | 2 | 0 | 0 |
| W12 | 2 | 0 | 0 |
| W1 | 3 | 0 | 0 |
| W2 | 3 | 1 | 0 |

| | | | |
|---|---|---|---|
| W3 | 3 | 0 | 0 |
| W4 | 3 | 1 | 1 |
| W5 | 3 | 1 | 1 |
| W6 | 3 | 0 | 0 |
| W7 | 3 | 0 | 0 |
| W8 | 3 | 0 | 0 |
| W9 | 3 | 0 | 0 |
| W10 | 3 | 0 | 0 |
| W11 | 3 | 0 | 0 |
| W12 | 3 | 0 | 0 |

Appendix 4. Survey Data *Isogenus nubecula*

| site | volume | pH | O2 | time | kick |
|---|---|---|---|---|---|
| W1 | 350 | 7.48 | 12.5 | 60 | possible |
| W2 | 200 | 7.53 | 11.9 | 60 | possible |
| W3 | 700 | 6.69 | 11.9 | 60 | possible |
| W4 | 1000 | 6.52 | 11.8 | 120 | possible |
| W5 | 750 | 7.83 | 11.4 | 45 | abs |
| W6 | 300 | 7.82 | 12.5 | 60 | abs |
| W7 | 750 | 7.67 | 11.6 | 90 | possible |
| W8 | 750 | 7.8 | 10.7 | 90 | possible |
| W9 | 750 | 6.75 | 11.8 | 90 | possible |
| W10 | 300 | 7.74 | 13 | 60 | possible |
| W11 | 500 | 6.63 | 11.6 | 45 | possible |
| W12 | 750 | 7.69 | 10.9 | 90 | possible |
